# Crosstalk between three CRISPR-Cas types enables primed type VI-A adaptation in *Listeria seeligeri*

**DOI:** 10.1101/2024.10.25.620265

**Authors:** Shally R. Margolis, Alexander J. Meeske

## Abstract

CRISPR-Cas systems confer adaptive immunity to their prokaryotic hosts through the process of adaptation, where sequences are captured from foreign nucleic acids and integrated as spacers in the CRISPR array, and thereby enable crRNA-guided interference against new threats. While the Cas1-2 integrase is critical for adaptation, it is absent from many CRISPR-Cas loci, rendering the mechanism of spacer acquisition unclear for these systems. Here we show that the RNA-targeting type VI-A CRISPR system of *Listeria seeligeri* acquires spacers from DNA substrates through the action of a promiscuous Cas1-2 integrase encoded by a co-occurring type II-C system, in a transcription-independent manner. We further demonstrate that the type II-C integration complex is strongly stimulated by preexisting spacers in a third CRISPR system (type I-B) which imperfectly match phage targets and prime type VI-A adaptation. Altogether, our results reveal an unprecedented degree of communication among CRISPR-Cas loci encoded by a single organism.

## Introduction

Clustered, regularly interspaced, short palindromic repeats (CRISPR) loci and associated (Cas) proteins are widespread prokaryotic adaptive immune systems that engage in RNA-guided surveillance and cleavage of foreign nucleic acids^1–5^. CRISPR loci are composed of short (∼30 bp) repetitive DNA elements with equally short “spacer” segments of foreign origin inserted between them. Naïve CRISPR immunity occurs in two phases. The first phase is adaptation, in which spacer sequences are captured from invading foreign genetic elements such as bacteriophages, or plasmids, and are inserted into the CRISPR locus to immunize the host against infection^1,6^. The second phase is interference, in which the CRISPR locus is transcribed and processed into small crRNAs, which associate with Cas nucleases to perform sequence-specific nucleic acid recognition and cleavage, and ultimately neutralize invading elements that match spacer sequences^2–5^. Two decades of bioinformatic mining have uncovered millions of unique CRISPR loci, categorized into 6 broad types and 35 subtypes with highly diverse gene content and function^7–9^.

Crucial to the adaptive nature of CRISPR immunity is the ability to acquire new spacers and thus gain heritable memory that protects the host from future infection. In all experimentally characterized CRISPR systems, spacer acquisition requires Cas1 and Cas2, which form a Cas1-2 integrase complex that captures sequences from free DNA ends, and inserts new spacer-repeat units into the CRISPR array, resulting in its expansion^10–13^. Phages readily escape CRISPR immunity by evolving mutations in the targeted genomic region that prevent recognition and/or cleavage by Cas nucleases^14^. To rapidly re-establish immunity against a changing viral population, many CRISPR systems have evolved mechanisms for “primed” adaptation. During this process, interference machinery equipped with crRNAs imperfectly matching a phage genome fails to directly neutralize the target, but strongly stimulates the acquisition of new spacers from target-proximal regions of DNA^15–17^. Priming has been best characterized in type I CRISPR systems, in which the Cascade surveillance complex bound to a mismatched target recruits the Cas1-2 integrase, which in turn recruits the helicase- nuclease Cas3^18^. The Cas1-2-3 complex then translocates along DNA emanating from the target, processing new spacers for immunization^18–20^. Primed adaptation has also been observed for type II CRISPR systems, which require the nuclease activity of Cas9 to generate free DNA ends that serve as substrates for further spacer acquisition^21^.

Many CRISPR-Cas systems do not encode Cas1-2, leading to the question of whether and how these systems are able to acquire new spacers^8^. Such systems are thought to adapt “in *trans*” using Cas1-2 machinery from other co-occurring CRISPR loci in the same genome. Adaptation in *trans* has been reported to accommodate spacer acquisition by the *Flavobacterium columnare* type VI-B system using co-resident type II- C Cas1-2, as well as the *Klebsiella pneumoniae* type IV-A3 system by type I-E adaptation machinery^22,23^. However, the generality of this phenomenon and the mechanisms underlying orthogonal array selection by a single Cas1-2 remain unclear.

Finally, the ability of some Cas1-2 machinery to install spacers in multiple distinct CRISPR loci raises the possibility that pre-existing spacers in one CRISPR locus might prime adaptation into other arrays, but this concept has not been experimentally investigated.

Here, we investigated spacer acquisition by the RNA-targeting type VI-A CRISPR system in its native host *Listeria seeligeri*. Type VI CRISPR systems use the nuclease Cas13 to engage mRNAs with complementarity to the spacer regions of crRNAs, and in response unleash nonspecific RNA degradation activity, leading to a state of cell dormancy that is incompatible with phage replication^24,25^. The *L. seeligeri* type VI-A system does not encode its own *cas1* or *cas2* genes, but our data indicate that it can acquire new spacers in *trans* using the Cas1-2 complex from a co-occurring type II-C system. Despite the RNA-targeting nature of type VI-A systems, we find that spacers are acquired from DNA in a transcription-independent manner, through a process that does not require the Cas13 nuclease. Flexibility in spacer acquisition is mediated by type II-C Cas1-2, which we find catalyzes promiscuous integration into type II and type VI arrays *in vitro*. Finally, we discovered that preexisting spacers in a third co-resident type I-B CRISPR locus strongly “cross-primes” type II-C Cas1-2 for spacer acquisition into the type VI-A locus. Our findings represent an unprecedented level of crosstalk between three CRISPR-Cas types in the same cell, raising the possibility that similarly complex interactions may govern CRISPR immunity in a wide range of microbes.

## Results

### Type II-C Cas1 and Cas2 mediate type VI-A spacer acquisition

We investigated the mechanism of spacer acquisition by the natural type VI-A CRISPR host *Listeria seeligeri* strain LS1^26^. Like all *Listeria* type VI-A systems, the LS1 type VI-A CRISPR locus lacks *cas1* and *cas2* genes (Figure 1A, S1A, Table S1).

**Figure 1.**
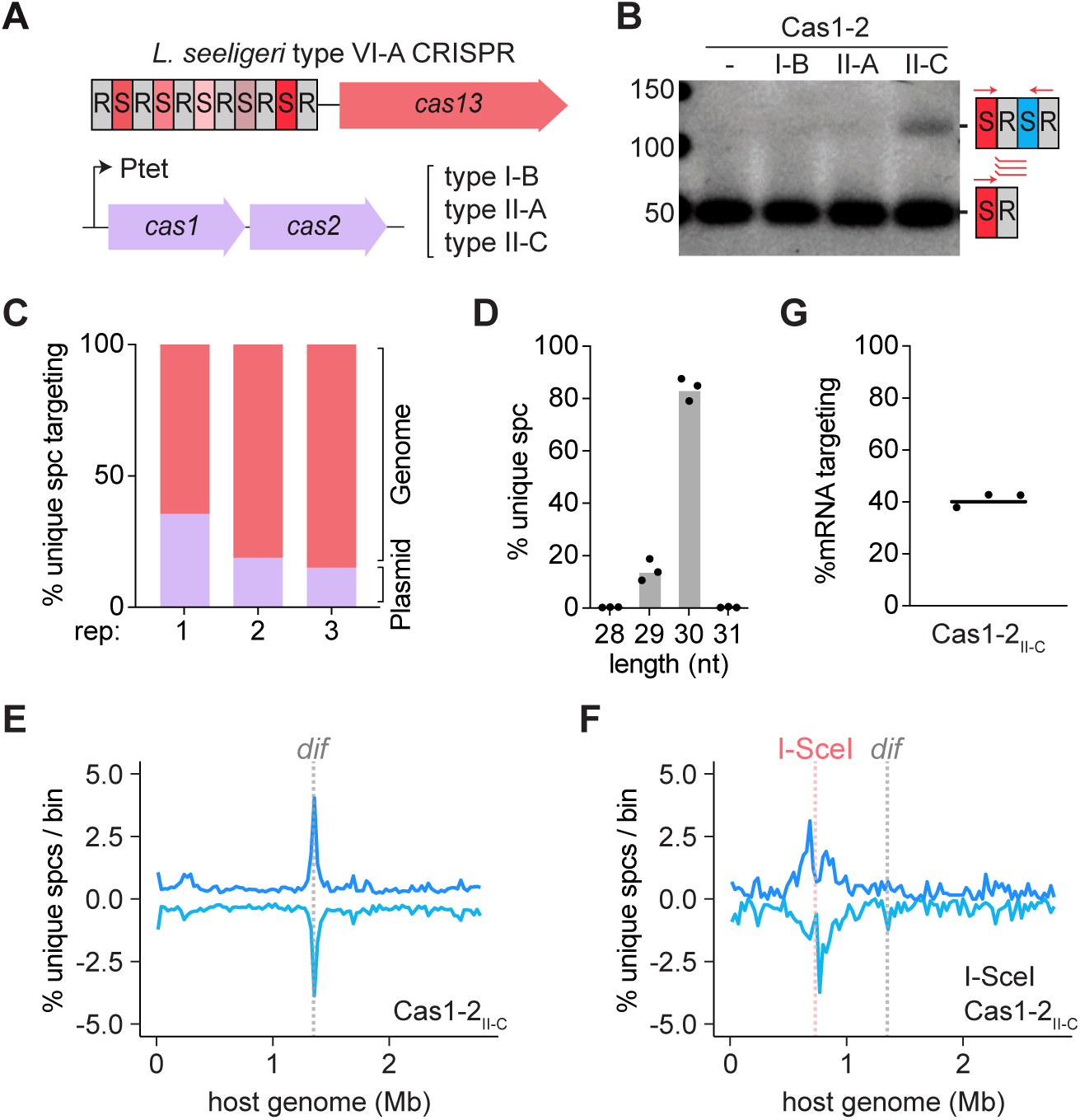
Type II-C Cas1-2 mediates type VI-A spacer acquisition. **(A)** Diagram of the experimental setup. Spacer acquisition was assayed in the native *L. seeligeri* type VI-A CRISPR locus during induction of different Cas1-Cas2 types from a plasmid. **(B)** Agarose gel showing the result of enrichment PCRs for new leader-adjacent spacers in the native type VI-A locus post Cas1-2 induction. **(C-G)** Deep sequencing analysis of type VI-A spacers acquired during type II-C Cas1-2 induction. **(C)** Percentage of unique spacers mapped to the plasmid or the *L. seeligeri* genome for each replicate (rep). **(D)** Length of newly acquired spacers. **(E)** Percentage of unique genome-mapped spacers acquired per 27976 bp bin (1% of the genome). Spacers were acquired from every genomic region but are particularly enriched near the *dif* site (gray line). The combination of unique spacers from n=3 replicates is shown. Positive values represent the top strand, while negative values represent the bottom strand. **(F)** Percentage of unique genome-mapped spacers acquired per bin after 24 hours of both type II-C Cas1-2 and I-SceI induction in a strain engineered with an I-SceI site (pink line). The combination of unique spacers from n=3 replicates is shown. **(G)** Percentage of unique genome targeting spacers that target mRNA.

However, of 111 *Listeria* genomes that harbor type VI-A loci, 93% encode one or more additional DNA-targeting CRISPR types that possess *cas1* and *cas2*, suggesting that adaptation in *trans* could mediate type VI-A spacer acquisition in these strains (Figure S1A, Table S1). For each of the three co-occurring CRISPR types (I-B, II-A, and II-C), we tested whether their cognate *cas1-2* alleles could stimulate the capture and integration of new spacers into the type VI-A array. In the case of type I-B CRISPR, we also co-expressed *cas4*, which is known to play roles in prespacer processing^27,28^. We induced *cas1-2* expression from a plasmid for 48 hours during 2 serial passages and subsequently performed enrichment PCR to preferentially amplify newly acquired spacers in the *cas13*-proximal position of the type VI-A CRISPR array. In these experiments, only expression of type II-C *cas1-2* was sufficient to robustly stimulate spacer acquisition in the native type VI-A array (Figure 1B).

We note that the *Listeria* type II-C CRISPR locus is variably identified as type II-A or type II-C. While the locus does encode a highly divergent homolog of the type II-A signature gene *csn2*, the Cas9 protein sequence is more closely related to well-characterized Cas9c proteins than to Cas9a (Figure S1B), and is much smaller than most Cas9a proteins. Furthermore, our results demonstrate that the *csn2*-like gene in this system is not strictly required for spacer acquisition. As type II-A systems originated from within the type II-C phylogeny, the *Listeria* system and its relatives may represent a recombinant hybrid^8^. Due to the clear distinctions between this locus and the canonical type II-A locus present in *Listeria*, in addition to the difference in results between the systems in this work and others^29–31^, we refer to it as a type II-C locus.

To analyze the distribution of genomic sequences sampled during type VI-A spacer acquisition, we performed deep sequencing of the expanded array amplicons and mapped the results to both the *cas1-2* expression plasmid and the *L. seeligeri* genome (Figure 1C, S2A-B). Most acquired spacers targeted the genome, but an outsized portion target the plasmid relative to its size (2.8 Mb genome vs. 5kb plasmid; Figure 1C). The vast majority of acquired spacers were 30 bp in length, with some shorter 29 bp spacers, consistent with the repertoire of native arrays (Figure 1D). To look for hotspots of acquisition, we split the *L. seeligeri* genome into 100 bins and counted the number of unique spacers targeting sequences in each bin (Figure 1E), which mitigates the impact of PCR bias. Spacers were acquired from most genomic regions, but there was a major enrichment of spacers targeting near the chromosomal terminus on both strands, as has been observed with type II-A spacer acquisition^12,13^. Specifically, the peak of spacers was centered at the 28 bp predicted *L. seeligeri dif* site, which differs from its experimentally verified equivalent in *Bacillus subtilis* by only a single nucleotide^32^. This enrichment is attributed to the double-stranded breaks (DSBs) that must occur at this site to resolve chromosomal dimers and concatemers prior to cell division. To determine whether DSBs are sufficient to cause type VI-A spacer acquisition, we engineered a unique 18 bp recognition site for the meganuclease I-SceI into the *L. seeligeri* genome, and induced expression of I-SceI along with type II-C Cas1-2 for 24 hours. The vast majority of spacers acquired in this experiment targeted regions centered around the cut site (Figure 1F, S2C), indicating that these DNA ends are preferred acquisition substrates.

Because type VI CRISPR systems target RNA, transcriptionally silent regions of the genome would not serve to produce functional spacers. We compared the newly acquired spacer repertoire to annotated open reading frames (ORFs) to see whether functional, mRNA-targeting spacers were preferentially acquired (Figure 1G). In all experiments, less than 50% of acquired spacers targeted mRNA, indicating that there is no mechanism for selection of functional type VI-A spacers at the level of acquisition.

We also considered that there could be some selection for functional spacers by acquiring preferentially from highly transcribed regions, as has been observed for type III-A CRISPR^33^. We compared the frequency of acquisition from each ORF to its read depth calculated from a previous transcriptomic analysis^25^, and found no correlation between transcript levels and spacer acquisition (Figure S3A). We also compared levels of transcription in 1 kb or 10 kb bins to acquisition in those bins and again found no correlation (Figure S3A). These data indicate that in *L. seeligeri*, type VI-A spacers are not selected for mRNA-targeting ability when acquired via type II-C adaptation machinery. Coupled with our observation of spacer acquisition from DNA ends, these results strongly suggest that DNA rather than mRNA is the primary substrate for type VI- A adaptation.

### Cas13 is not required for type VI-A spacer acquisition

How is the type II-C Cas1-2 complex recruited to integrate spacers into an orthogonal type VI-A array? As Cas13 is the only ORF in the type VI-A locus, we wondered whether Cas13 could play a role in spacer acquisition. We repeated the type II-C Cas1-2 induction experiments in strain backgrounds where either the nuclease activity of Cas13 was removed by inactivation of both HEPN domains (dCas13), or where an early stop codon was introduced to make a Cas13 null (Cas13_Y80*_). These Cas13 mutants were confirmed to be nonfunctional in target plasmid interference (Figure S4A). The absence of Cas13 had no detectable effect on the substrate source, length, distribution, or mRNA-targeting frequency of acquired spacers (Figure 2A-D, S3B-C, S4B-C), indicating that Cas13 is neither required for nor influences type VI-A spacer acquisition. We were surprised that the presence of Cas13 had no impact on the frequency of mRNA-targeting spacers acquired (Figure 1G vs. 2B). In principle, Cas13 should be activated by expressed target mRNAs, causing the cell to enter dormancy, and this effect should be abrogated with non-functional Cas13 alleles (dCas13 or Cas13_Y80*_). These results could arise from insufficient target expression levels to trigger Cas13 activity^34^; detection of spacers originating from dormant yet intact cells in the culture; and/or inactivation of Cas13 in cells acquiring self-targeting spacers. However, we are unable to distinguish between these possibilities in bulk assays.

**Figure 2.**
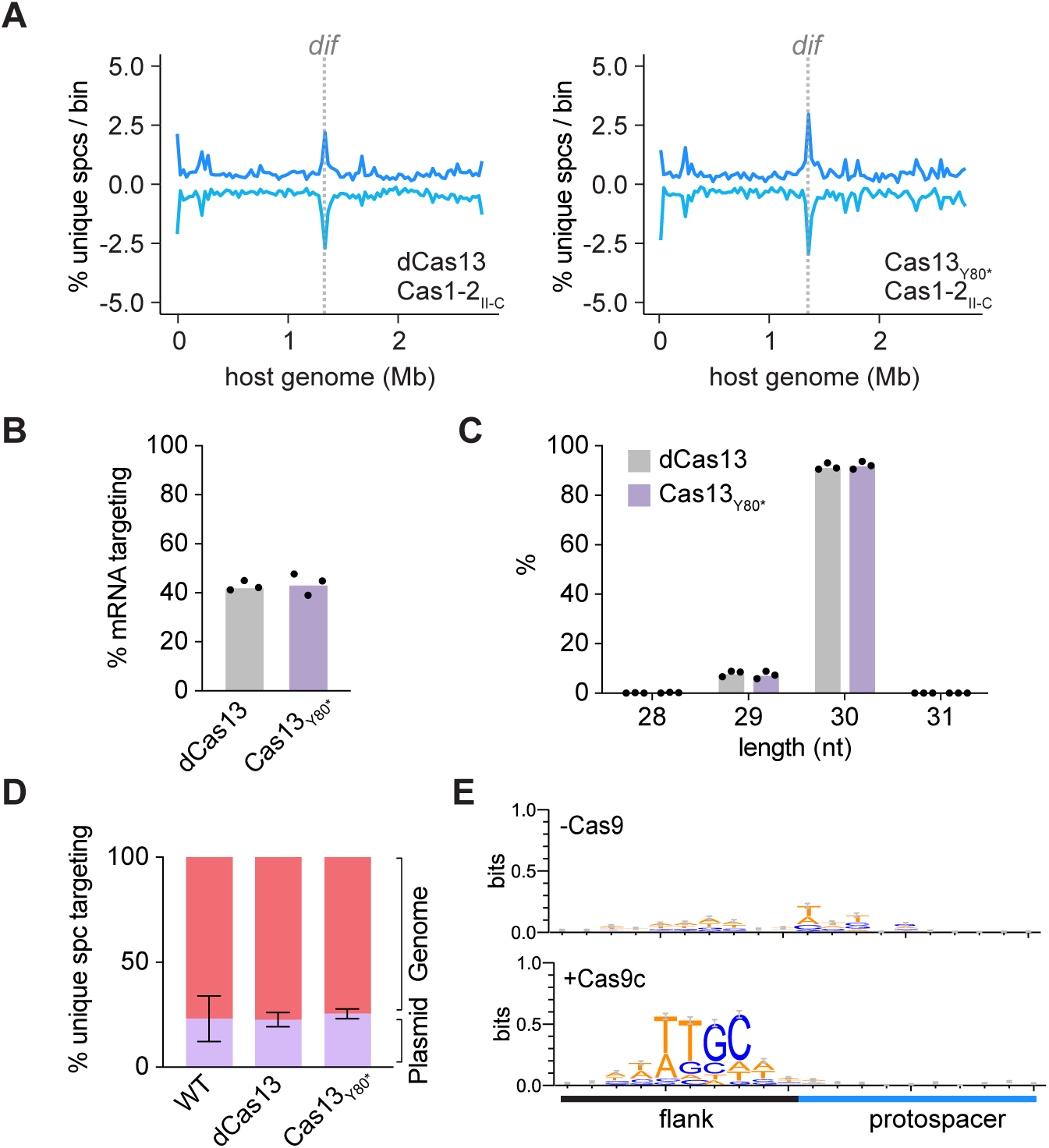
Contribution of Cas nucleases to type VI-A spacer acquisition. **(A)** Percentage of unique, genome-mapped, type VI-A spacers acquired per bin during type II-C Cas1-2 induction in dCas13 (left) or Cas13_Y80*_ (right) mutants. The combination of unique spacers from n=3 replicates is shown. Positive values represent the top strand, while negative values represent the bottom strand. (**B-D)** Analysis of mRNA targeting ability (B), length (C), and plasmid vs. genomic targets of newly acquired spacers from the experiment in (A). In (D), the data for the WT condition are the same as in Figure 1C. **(E)** Sequence motif resulting from alignments of of genome-mapped type VI-A spacer targets acquired during type II-C Cas1-2 expression, in the presence or absence of Cas9c. Each plot represents a combination of two biological replicates.

The *cas1-2* genes we expressed are derived from a type II-C CRISPR locus that also encodes Cas9c, which we reasoned could play a role in type VI-A spacer acquisition. We were unable to do experiments in a strain that natively contains both the type VI-A and type II-C systems as these strains contain anti-CRISPR activity from an unidentified source that renders the type II-C system inactive^31^. We instead inserted the *cas9c* gene and its cognate tracrRNA into the LS1 chromosome along with type II-C arrays containing 1 or 10 spacers, and repeated the type II-C Cas1-2 induction experiments. Under these conditions, type VI-A spacers were still acquired with similar patterns to those observed in the absence of *cas9* (Figure S4D), but the preponderance of newly acquired spacers matched target sequences flanked by a protospacer adjacent motif (PAM) identical to the reverse complement of natural type II-C targets (3’ NNGGCA; Figure 2E). These data suggest that Cas9c participates in the type VI-A spacer acquisition process and is responsible for specifying a type II-C PAM. Our results extend the previous finding that type II-A Cas9 mediates the selection of targets with the correct PAM during adaptation^35^, and are consistent with observations of type II-C PAMs during type VI-B spacer acquisition in *F. columnare*^22^. Altogether, these data indicate that in a strain with type II-C and VI-A CRISPR-Cas systems, the spacers acquired in the type VI-A locus follow the patterns of type II-C spacer acquisition.

### Type II-C Cas1-2 integrates prespacers into type II and type VI arrays in vitro

Since Cas13 is not required to recruit the type II-C Cas1-2 complex to the type VI-A array, we reasoned that sequence similarities between the two arrays may allow either to be recognized by Cas1-2 for insertion of new spacers. The two arrays contain 36 bp direct repeat sequences sharing 54% nucleotide identity, spacers averaging 30 bp in length, and exhibit some similarity in the leader sequence, especially in the array- proximal region which is important for directing the polarity of spacer acquisition in type II-A systems^36^ (Figure 3A). To test whether type II-C Cas1-2 exhibits a preference for these arrays, we first separately expressed and purified the *L. seeligeri* type II-C Cas1 and Cas2 proteins from *E. coli* (Figure S5A). Next, we tested their ability to perform half-site integration into plasmids containing single-spacer type VI-A, II-A, II-C, and I-B arrays *in vitro* when supplied with dsDNA prespacer fragments containing 4 nt ssDNA overhangs on both ends^37^ (Figure 3B). Half-site integration was assayed using one primer downstream of the array on the plasmid substrate, and one primer matching the prespacer, such that preferential integration anywhere on the plasmid could be detected. Cas1-2 catalyzed half-site integration only into plasmids containing the type II-A, II-C, and VI-A arrays, and this occurred at the expected leader-repeat junction (Figure 3C; products were also confirmed by sequencing). Next, we tested whether the leaders and/or repeats were required for integration by separately deleting the leader or repeat-spacer-repeat from plasmid substrates. Half site integration was abrogated in both L- and RSR-only plasmids for both type II-C and type VI-A (Figure 3D), indicating that both the leader and repeats are required for prespacer integration in vitro. These experiments indicate that the *L. seeligeri* type II-C integration machinery exhibits inherent promiscuity with respect to array recognition, and is sufficient to integrate dsDNA prespacers into type VI-A CRISPR arrays.

**Figure 3.**
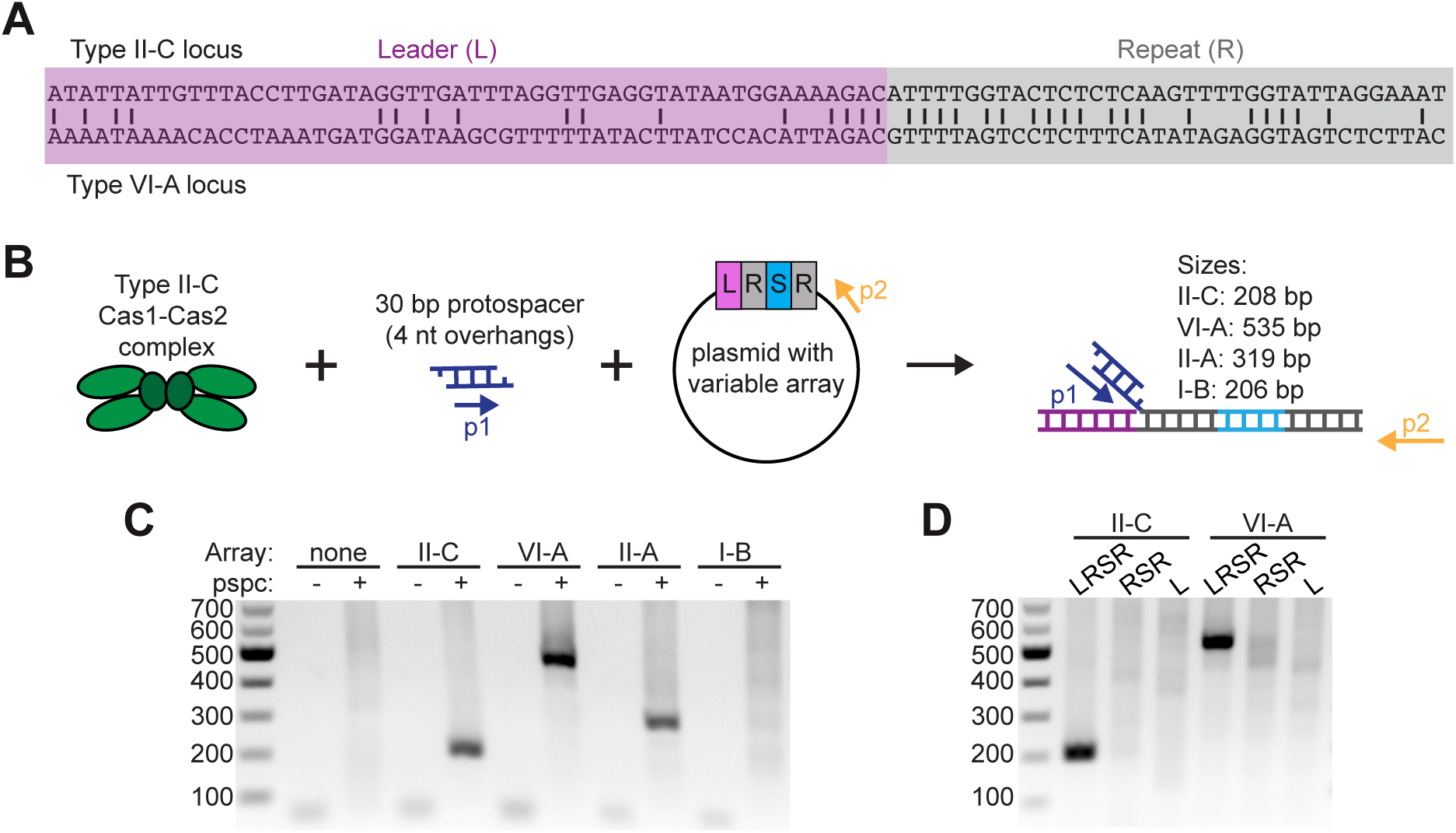
Type II-C Cas1-2 mediates half-site integration into type II and type VI-A arrays in vitro. **(A)** Comparison of the *L. seeligeri* type II-C and type VI-A leader and direct repeat sequences. **(B)** Schematic of the assay to detect half-site integration by type II-C Cas1-2 in vitro. Purified Cas1-2 protein was incubated with a 30 bp prespacer (with 4 bp overhangs; pspc) and a plasmid with variable CRISPR arrays. The location of primers p1 and p2, as well as the expected sizes of the PCR product from leader-adjacent half-site integration into different arrays, are noted. **(C)** Integration assay PCR on indicated samples. Sizes are consistent with half-site integration at the leader adjacent repeat for type II-C, type VI-A, and type II-A arrays. **(D)** Integration assay PCR with indicated type II-C or VI-A plasmids. LRSR, Leader-Repeat-Spacer-Repeat. RSR, array only. L, Leader only. Results are representative of n=3 experiments. Samples without pspc were incubated with Cas1-2 and plasmid only. No array control contains the parental vector.

### Acquisition of type VI-A spacers during phage infection

We wondered whether type II-C Cas1-2 expression would enable acquisition of protective type VI-A phage-targeting spacers during infection. We infected *L. seeligeri* cells expressing type II-C Cas1-2 with the temperate listeriaphage U153 at an MOI of 0.1 and isolated genomic DNA 48 hours post infection (hpi) from surviving cells. Most cells had not acquired a type VI-A spacer, indicating there are other ways for the cells to gain resistance to U153. Regardless, we deep sequenced the expanded type VI-A arrays from each sample and mapped acquired spacers to both the host and phage genomes. A disproportionate number of the type VI-A spacers detected in this experiment targeted the phage genome (Figure 4A; 2.8 Mb host genome vs. 42 kb phage genome), indicating that type II-C Cas1-2 can catalyze type VI-A adaptation to infecting phage, with no clear strand bias or subgenomic sites of enrichment (Figure 4B, S6B). Interestingly, a new spacer acquisition hotspot appeared in the host chromosome during U153 infection. This region corresponded to host genomic regions adjacent to the U153 prophage integration site, which is located within the *comK* gene^38^ (Figure 4C, S6A). These results raised the possibility that prophage integration stimulates spacer acquisition, as it is a process that involves DSB formation^39^. To test this, we constructed a lytic U153 phage mutant (U153_lyt_) lacking the lysogenic operon and repeated the infection and acquisition experiments. Spacers targeting the lytic mutant were acquired at the same rate as WT phage, but we no longer observed strong enrichment of spacers originating from near the prophage integration site (Figure 4A,D,E, S6C,D), demonstrating that type VI-A spacers can be acquired during lytic infection. To further explore the relationship between lysogeny and type VI-A adaptation, we isolated U153 lysogens, induced type II-C Cas1-2, and monitored acquisition of new spacers in the type VI-A array. Under these conditions, we observed a dominant peak of newly acquired type VI-A spacers targeting the prophage and surrounding regions of the host genome (Figure 4F, S6E), which was completely dependent on type II-C Cas1-2 expression (Figure S6F). We reasoned that spacer acquisition from the prophage could be explained by spontaneous prophage reactivation by subpopulation of the cells in the culture, followed by prophage excision and subsequent phage genome injection into neighboring lysogens. To investigate this, we deleted the integrase gene from the prophage, which impaired its ability to produce viable PFU during by 5 orders of magnitude during prophage induction (Figure S6G). Surprisingly, when we repeated the type II-C Cas1-2 induction experiments in the lysogen harboring Δ*int* prophage, we observed an equally strong enrichment of type VI-A spacers targeting the prophage and surrounding regions (Figure 4G, S6H). That spacers were acquired from this inert “locked-in” prophage mutant suggests that some property of the U153 sequence itself, rather than the phage life cycle, strongly stimulates spacer acquisition.

**Figure 4.**
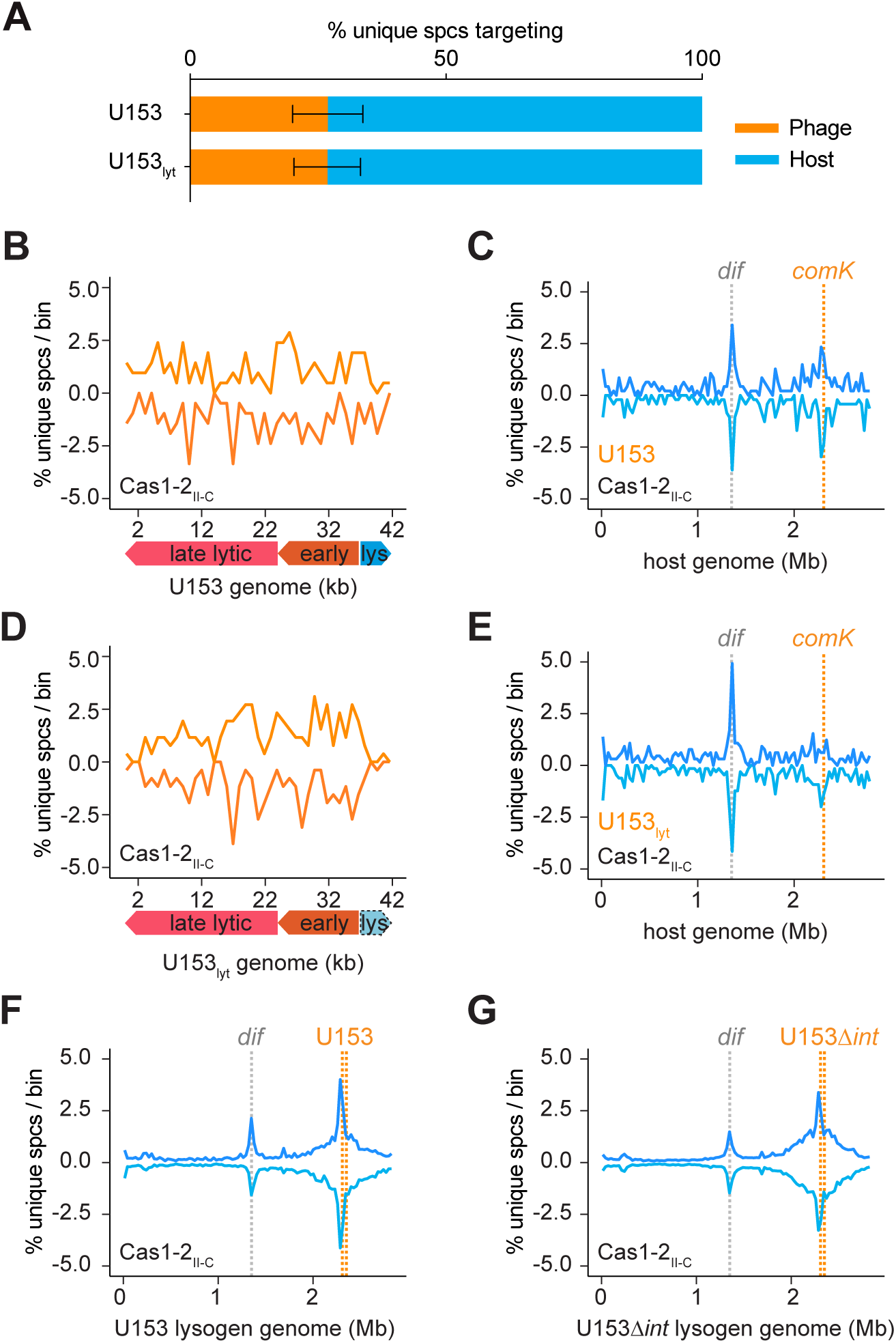
Acquisition of phage-targeting type VI-A spacers during type II-C Cas1-2 induction. **(A)** Percentage of unique acquired spacers targeting the host or phage genome during infection with U153 phage or strictly lytic mutant (U153_lyt_) in cells expressing type II-C Cas1-2. **(B-E)** Percentage of unique spacers per bin from the experiment described in A, mapped to the phage (**B,D**) and host (**C,E**) genomes. The combination of unique spacers from n=3 replicates is shown. In B and D, the bin size is 986 bp; in C and E, the bin size is 1% of the host genome. Prophage integration site *comK* (orange) and *dif* site (gray) are indicated. **(B-C)** Spacers acquired during WT U153 infection, where some infection events may result in prophage integration. **(D-E)** Spacers acquired during infection with U153_lyt_ which lacks integration machinery. **(F-G)** Percentage of unique lysogen genome-mapped type VI-A spacers acquired per bin (1% of the lysogen genome) during type II-C Cas1-2 induction. The combination of unique spacers from n=3 replicates is shown. Boundaries of the integrated prophage shown (orange). **(F)** Spacers acquired in the WT U153 lysogen, where the prophage can potentially reactivate. **(G)** Spacers acquired in the U153Δ*int* lysogen, in which the U153 integrase gene has been knocked out of the lysogen genome, largely removing the prophage’s ability to excise.

### Type I-B CRISPR primes type II-C Cas1-2 for type VI-A spacer acquisition

The experimental strain of *L. seeligeri* we used for our assays (LS1) contains, in addition to a native type VI-A locus, a type I-B CRISPR-Cas locus that is largely transcriptionally silent under laboratory growth conditions (Figure S7A), and does not interfere against plasmids containing matching protospacers (Figure S7B). However, the type I-B system is functional if expressed under the control of an inducible promoter^31^.

Upon further analysis we noticed that this type I-B locus contains one natural spacer that targets the U153 genome with a single mismatch (Figure 5A). This native type I-B locus does not offer any protection during U153 infection (Figure 5B), as WT LS1 and a mutant lacking the entire type I-B CRISPR-Cas locus are equally permissive to phage infection. We hypothesized that in a subpopulation of cells, or at some low level, the type I-B system is expressed and able to prime type VI-A spacer acquisition through type II-C Cas1-2. To explore this possibility, we lysogenized mutants lacking either the type I-B helicase-nuclease Cas3 or the entire type I-B CRISPR-Cas locus with U153 and repeated the spacer acquisition experiments with induction of type II-C Cas1-2. In both deletion mutants, spacers targeting the prophage and surrounding genome were no longer preferentially acquired in the type VI-A array (Figure 5C-E, S8A-B,E). We also found that phage-targeting type VI-A spacers were still acquired during U153 infection of the Δtype I-B mutant, indicating that type I-B priming is not strictly required for type VI-A adaptation to phage (Figure S8C). To test whether the endogenous type I-B Cas1 and Cas2 are involved in type VI adaptation, we deleted both from the LS1 genome, which had no impact on type VI-A spacer acquisition, further confirming that type II-C Cas1-2 are solely responsible for spacer integration (Figure 5F, S8D). Finally, to test whether a type I-B spacer is sufficient to prime type VI-A acquisition by type II-C CRISPR, we ectopically integrated a type I-B spacer targeting the host genome with a single mismatch under the native type I-B array promoter, and repeated the type II-C Cas1-2 induction experiments. Strikingly, we observed the emergence of a new hotspot of type VI-A spacers mapping near the engineered type I-B target site (Figure 5G, S8F), which was dependent on the presence of Cas3 (Figure 5H, S8G) and the native type I-B CRISPR-Cas system (Figure S8H). Altogether, these data indicate that type I-B CRISPR can prime the type II-C Cas1-2 integrase to capture and integrate new spacers into the type VI-A CRISPR array.

**Figure 5.**
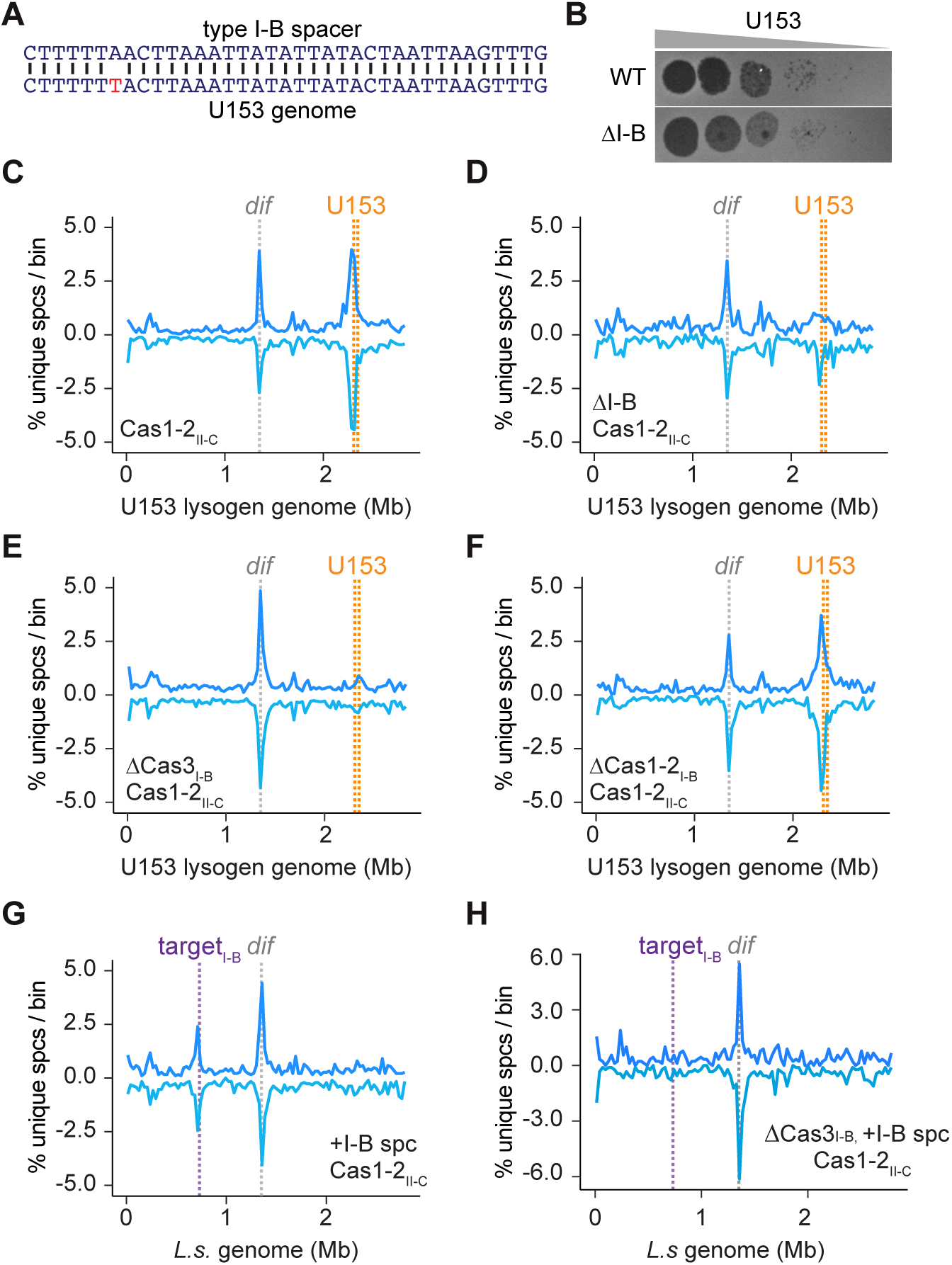
Type I-B CRISPR primes type II-C Cas1-2 for type VI-A spacer acquisition. **(A)** Comparison of a type I-B spacer that exists in the LS1 genome to a sequence in the U153 genome. A single mismatch is highlighted in red. **(B)** U153 infection of lawns of WT LS1 or LS1 lacking the entire type I-B locus (ΔI-B). Representative of n=3 experiments. **(C-F)** Percentage of unique lysogen-genome-mapped type VI-A spacers acquired per 1% bin during type II-C Cas1-2 induction. The combination of unique spacers from n=2 replicates is shown. Boundaries of the integrated prophage (orange), and *dif* site (gray) shown. **(C)** Spacers acquired in WT U153 lysogens; as in Fig. 4F, but from 2 distinct biological replicates completed alongside the experiments in D and E. **(D)** Spacers acquired in ΔI-B U153 lysogens. **(E)** Spacers acquired in U153 lysogens lacking the Cas3 gene in the type I-B locus. **(F)** Spacers acquired in U153 lysogens lacking the Cas1 and Cas2 genes in the type I-B locus. (**G-H)** Percentage of unique genome-mapped spacers acquired per 1% bin during type II-C Cas1-2 induction in the presence of a type I-B spacer targeting a genomic location (purple) with a single mismatch. The combination of unique spacers from n=2 replicates is shown. **(G)** Spacers acquired in WT LS1. **(H)** Spacers acquired in ΔCas3_I-B_ LS1.

## Discussion

In this study, we sought to understand how a type VI-A CRISPR system can acquire new spacers without encoding its own Cas1 and Cas2 machinery. We found that in *Listeria seeligeri*, type II-C Cas1 and Cas2 can integrate new spacers into the native type VI-A locus. Type VI CRISPR acquisition using the type II-C integrase complex has previously been reported in the distantly related type VI-B system of *Flavobacterium columnare,* where the type II-C Cas1 and Cas2 are only 33% and 37% identical to the *L. seeligeri* system, respectively. This suggests that the promiscuity of type II-C Cas1-2 enzymes, or of type VI arrays, may be a widespread phenomenon.

During this study, we also serendipitously discovered a role for type I-B priming in type VI-A spacer acquisition by using a phage that has a single mismatch to a spacer natively present in the *L. seeligeri* type I-B array. While this spacer does not provide broad protection during infection (Figure 5B), it is able to prime type II-C Cas1-2 to capture spacers near the mismatched target and integrate them into the type VI-A array. During type I priming, Cascade recruits type I Cas1-2 to mismatched targets^18^; perhaps it is able to physically interact with divergent type II-C Cas1-2 machinery as well.

Alternatively, target recognition by the priming spacer could stimulate a low level of target DNA cleavage by Cas3, which would supply linear DNA ends to serve as spacer acquisition substrates. The extent of such “cross-priming” between distinct CRISPR types in the same genome remains an outstanding question. This phenomenon requires either Cas1-2 machinery that can act in *trans* on multiple CRISPR arrays or communication between the priming CRISPR system and multiple distinct Cas1-2 complexes. Here we have demonstrated that both mechanisms exist, and could provide a path for rapid dissemination of established immunological memories across CRISPR-Cas loci. As functional PAM sequences are selected during the spacer acquisition process, relatively PAM-flexible type III and type VI CRISPR systems are well-poised to receive spacers from orthogonal Cas1-2 machinery acting in *trans*. Cross-priming might be particularly advantageous in the event of infection by a phage refractory to targeting by the priming CRISPR type, due to expression of anti-CRISPRs or other resistance mechanisms. Indeed, it has also been shown that some type III CRISPR systems can mediate interference using type I crRNAs, which enables robust protection against phages containing target site mutations^40^. Taken together, these and our findings indicate that distinct co-resident CRISPR types can influence each other at all levels of immunity.

It is worth noting that during cross-priming in the U153 lysogen, all acquired spacers are self-targeting, despite the priming spacer targeting a phage-derived sequence. In DNA targeting CRISPR systems, this would not be advantageous, as newly acquired prophage-matching spacers could then be used to target and cleave the host genome. However, type VI CRISPR systems sense RNA, and therefore the target must be transcribed to trigger Cas13 activity. Prophages do not express the majority of their genes, as these make proteins involved in replication and lysis. However, upon reactivation, these lytic mRNAs are again produced. Thus, if a type VI spacer targeting a lytic mRNA of the prophage were acquired through the priming mechanism discovered here, this immune system would be “off” until the phage reactivates. This mechanism could potentially allow a cell to maintain beneficial genes present in a prophage but stop it from reactivating and lysing the cell.

Why do so few type VI CRISPR systems encode Cas1 and Cas2 genes? We propose that it may be evolutionarily advantageous to couple acquisition of type VI spacers with presence of a DNA-targeting immune system, such as type I or II CRISPR. Type VI CRISPR activation leads to cell dormancy, and as long as the activating mRNA target is present, the cell will not escape dormancy. However, we have previously shown that in the presence of restriction-modification systems, cells that have undergone Cas13-mediated dormancy can be resuscitated due to destruction of the phage DNA that is transcribed to produce the activating mRNA^41^. Perhaps using Cas1-2 from a DNA targeting system gives cells with type VI immunity a better chance to exit dormancy if this machinery is also used to acquire DNA-targeting spacers from the same target. Further studies will be needed to shed light on the advantages and disadvantages of harboring multiple CRISPR systems that can interact with each other.

## Acknowledgments

We thank Marshall Godsil for finishing construction of U153-cmR, and for especially helpful suggestions. We also thank Monica Guo for help with protein purification, and Albina Kozlova, JC Alexander, and David Brinkley for previous work to detect type VI-A spacer acquisition. We would also like to thank all members of the Meeske lab for helpful discussion. Work in the Meeske lab is supported by NIGMS (R35GM142460), NSF (FAIN2235762), and the University of Washington Royalty Research Fund. SRM is a Jane Coffin Childs Memorial Foundation Postdoctoral Fellow. AJM is a Rita Allen Foundation Scholar.

## Author Contributions

The study was conceived by SRM and AJM. SRM performed all experiments described in the paper. SRM and AJM wrote and edited the paper.

## Supplemental Information

Figures S1-S8.

Table S1, Bioinformatic analysis of CRISPR content in *Listeria spp*. genomes.

Table S2, Strains, plasmids, and oligonucleotide primers used in this study.

## Methods

### Bacterial strains and growth conditions

All *Listeria seeligeri* strains were cultured in Brain Heart Infusion (BHI) broth or agar at 30 °C, and all *Escherichia coli* strains were grown in lysogeny broth (LB) or agar at 37°C supplemented with appropriate antibiotics (specified below). Plasmids were initially cloned in DH5α or Turbo Competent *E. coli* (New England Biolabs) and miniprepped.

For introduction into *L. seeligeri*, plasmids were then transformed into a conjugative donor strain of *E. coli* (specified below).

### Plasmid construction

Broadly, plasmids were constructed by Gibson assembly of PCR purified inserts with restriction digested plasmids, except in the case of spacer cloning, where annealed oligos were directly ligated into cut vectors. Details of plasmid construction, along with all plasmids and oligonucleotides used in this study can be found in Table S2.

### E. coli–L. seeligeri conjugation

Donor *E. coli* strains S-17 λpir, SM10 λpir, or β2163 Δ*dapA* (for allelic exchange) carrying *E. coli*–*Listeria* shuttle vectors were cultured overnight in LB medium with the appropriate antibiotic: 25 µg/ml chloramphenicol (for pPL2e-derived plasmids), 50 µg/ml kanamycin (for pAM326-derived plasmids), or 100 µg/ml ampicillin (for pAM8-derived plasmids). Recipient *L. seeligeri* strains were grown overnight in BHI medium supplemented with 1 µg/ml erythromycin (for pPL2e-derived plasmids), 10 µg/ml chloramphenicol (for marked lysogens or pAM8-derived plasmids), or 50 µg/ml kanamycin (for pAM326-derived plasmids) at 30°C. The saturated donor and recipient cultures (100 µl each) were combined in 10 ml BHI and concentrated onto a 0.45-µm membrane filter disc (Millipore-Sigma) using vacuum filtration. The filter was then placed on BHI agar containing 8 µg/ml oxacillin, which weakens the cell wall, enhancing conjugation, and incubated at 37°C for 4 hours. After incubation, the cells were resuspended in 2 ml BHI and plated on selective BHI medium containing 50 µg/ml nalidixic acid, which kills donor E. coli but not recipient Listeria, along with the appropriate antibiotic for plasmid selection. Transconjugants were isolated after 2–3 days of incubation at 30°C.

### Cas1-2 induction experiments

*L. seeligeri* strains were grown overnight in BHI with the appropriate antibiotics. The next day, cultures were diluted 1:1000 into BHI with antibiotics and 100 ng/mL anhydrotetracycline (aTc) for induction of the Ptet promoter. After 24 hours, cultures were similarly passaged once more into fresh media with aTc and after another 24 hours (48 hours total induction) cells were pelleted by centrifugation and frozen at -20°C prior to genomic DNA (gDNA) extraction.

### Sample preparation and array amplification for next-generation sequencing

Cell pellets were resuspended in PBS and lysed by incubation with 2mg/mL lysozyme and 1% N-lauroylsarcosine. Genomic DNA was isolated by phenol/chloroform extraction followed by ethanol precipitation. 100ng of genomic DNA were used as input for the type VI-A array enrichment PCR with Q5 High-Fidelity DNA polymerase (NEB) and 0.4µM final concentration of each primer: oDB026 (binds to pre-existing leader-adjacent spacer (spc1)), and oSM039, oSM040, oSM041 (each of which binds to the type VI-A repeat with the final two 3’ nucleotides differing to match any nucleotide except those of spc1). Reactions were performed with a 57.5°C primer annealing temperature and 10 second extension time, and then run on a 2% agarose gel from which bands larger than 100bp were extracted using a QIAquick Gel Extraction Kit. Amplicons were prepared for sequencing with the TrueSeq Nano DNA Library Prep protocol (Illumina) and sequenced with the NextSeq or NovaSeq X Plus platforms (Illumina).

### High throughput sequencing data analysis

Newly acquired spacer sequences, defined as those flanked by spc1 and direct repeat on one side, and a second repeat on the other side, were extracted from the raw Illumina FASTQ files. Instances of each spacer sequence were counted and unique spacers were mapped to host, plasmid, or phage genomes using bowtie2 with default parameters. Sequences mapping to the existing type VI-A array were assumed to be PCR artifacts and discarded (for samples with few acquired spacers, these were the majority of reads). Next, unique mapped spacers with over 10 instances were tallied for defined bins in each genome. All spacers derived from the same strand and alignment coordinates were considered to be the same unique spacer. All plots were made in R, and replicates were combined for plotting as indicated in the figure legends. Plots with full counts of all newly acquired spacers mapped to the genome for each replicate can be found in supplemental figures. For determining mRNA targeting ability of newly acquired spacers, spacers were compared to the strand and orientation of LS1 open reading frames. To test for correlation with transcription, spacer abundance per bin was compared to transcription levels of previously published RNA-seq data^25^ using a Spearman’s rank correlation test. To generate sequence motifs, the 30 bp regions flanking all 30 bp spacer sequences that mapped to the LS1 genome were extracted and analyzed using WebLogo.

### Gene deletions

Allelic replacement in *L. seeligeri* was performed as previously described^42^. Briefly, 500- 1000 bp homologous sequences flanking the region targeted for replacement were cloned into the suicide plasmid pAM215, which contains lacZ and chloramphenicol resistance (cat) markers. The plasmid was introduced into *L. seeligeri* strains via conjugation as outlined above, and 100 µl of the resuspended cells was plated on selective media containing 50 µg/ml nalidixic acid and 15 µg/ml chloramphenicol.

Transconjugants (integrants) were isolated and passaged (grown to saturation, then diluted 1,000-fold) 3 times in BHI without antibiotics. The passaged cultures were then diluted and plated on BHI plates with 100 µg/ml 5-bromo-4-chloro-3-indolyl β-d-galactopyranoside (X-gal), followed by incubation for 2–3 days at 30°C. White (lacZ–) colonies (excisants) were selected, confirmed to be chloramphenicol-sensitive, and the deletion was verified by PCR with primers flanking the target region.

### Cas1 and Cas2 purification

Codon optimized *L. seeligeri* type II-C Cas1 and Cas2 were cloned into pKS22b-His-SUMO, and the resulting plasmids were separately transformed into BL21(DE3) *E. coli*. Cells were grown in 1 L of LB with 100 μg/ml carbenicillin shaking at 37°C until reaching an OD_600_ of 0.6, and then induced overnight at 18°C with 1 mM isopropyl-β-D-thiogalactopyranoside (IPTG). The cells were pelleted, resuspended in lysis buffer (50 mM Tris-HCl, pH 7.5, 500 mM NaCl, 20 mM imidazole, 10% glycerol, 1mM DTT, 1mg/ml lysozyme, 1mM PMSF, 15 units DNAseI, 0.5mM MgCl2), and incubated on ice for one hour before sonication. Sonicated lysates were centrifuged, and the cleared lysates were passed over Ni-NTA agarose resin (MilliporeSigma) at 4°C.The resin was washed with wash buffer (50 mM Tris-HCl, pH 7.5, 500 mM NaCl, 20 mM imidazole, 10% glycerol, 1mM DTT) and the protein eluted in wash buffer supplemented with 400 mM imidazole. Eluted protein was dialyzed overnight in fresh wash buffer with Ulp1-His protease to remove the His-SUMO tag. The next day, the protein was again passed over Ni-NTA resin to remove both the protease and any remaining uncleaved protein.

The flowthrough was collected, concentrated, and further purified on a Superdex -75 Increase 10/300 in protein buffer (50 mM Tris-HCl, pH 7.5, 500 mM NaCl, 10% glycerol, 1mM DTT). Fractions containing the protein of interest were combined, concentrated, aliquoted, and stored at -80°C.

### In vitro integration assays

Oligos oSM97/oSM98^37^ were hybridized in integration buffer (20mM Tris-HCl pH 8, 25 mM NaCl, 10 mM MgCl_2_, 1mM DTT, 10% DMSO) by heating to 95°C and slowly cooling to room temperature to create a 30bp protospacer with 4bp overhangs. 5 μM Cas1 and Cas2 were combined and incubated on ice for 30 minutes, then diluted to 47 nM and incubated with 200 nM protospacer on ice for 15 minutes. 20 nM of target plasmids that had been purified from DH5α *E. coli* were then added and incubated at 37°C for 30 minutes, then quenched with 0.4% SDS and 50 mM EDTA. Plasmids were purified using the DNA Clean & Concentrator-5 kit (Zymo), eluted in 8 µl of water, and 1 µl of 1:6 diluted plasmid was used as input for half-site integration PCRs. PCRs were performed using Q5 Hot Start High-Fidelity 2x Master Mix (NEB) per the manufacturer’s instructions with a 30 second extension time using primers oSM097 (p1 in Figure 3) and oSM029 (p2 in Figure 3). Reaction products were analyzed on a 1.5% agarose gel containing ethidium bromide. PCR products were sequenced to confirm that bands corresponded to the expected leader-adjacent half-site integration.

### Phage propagation and infections

All phage infections were performed in BHI with 5 mM CaCl_2_. Top agar lawns were made by combining 100 µL of overnight culture with CaCl_2_ and 5 mL of molten BHI agar. Phage stocks were generated from single plaques formed on top agar lawns of LS1 ΔRM1 ΔRM2 ΔCRISPR_VI-A_. In the case of U153_lyt_, cells also contained pAM324, which expresses Cas9 targeting the phage integrase, allowing for isolation of a spontaneous phage mutant that lost the entire lysogenic operon (confirmed by PCR with oSM101 and oSM288). Phage stocks were made by infecting 5 mL of *L. seeligeri* at OD_600_ of 0.1 with the plaque, allowing infection to proceed overnight, and filtering the resulting lysate through a 0.45-μm-pore syringe filter. The stock was titered on top agar lawns to identify the plaque forming units (PFU) per µL, and this information was used to infect experimental strains at a multiplicity of infection of 0.1. Infection and addition of atc to induce Cas1-2 occurred at the same time. After 48 hours of infection, cells were pelleted by centrifugation and stored at -20°C prior to gDNA extraction. To determine PFU production by lysogens, cultures at OD_600_ of 0.1 were treated with 2 µg/mL mitomycin C overnight, filtered and tittered as above.

### U153-cmR construction

A constitutively expressed *cat* gene was inserted into a non-essential region of the U153 genome, downstream of the lysin gene. Briefly, the kanR plasmid pAM591 was constructed containing the *cat* gene flanked by 500 bp of U153 genomic sequence surrounding the desired insertion site. After introducing pAM591 into LS1 via conjugation, the resultant strain was infected with phage U153 to enable recombination with pAM591. Infection was performed at an MOI of 0.1, proceeded overnight, and a phage stock was harvested as above. This stock represented a heterogeneous mixture of wild-type and recombinant phage, which was then used to infect a fresh LS1 culture at an MOI of 1 for 1 hour. Lysogens were then selected by plating the infected cells on BHI + chloramphenicol. Chloramphenicol-resistant lysogens were confirmed to be kanamycin-sensitive, then the recombinant prophage was induced by treatment with 2 µg/mL mitomycin C, and plaques were isolated on a lawn of wild-type LS1 infected with the induced culture filtrate. A stock of recombinant U153-cmR was prepared by expanding a single plaque as above.

### Bioinformatic analysis of CRISPR loci in Listeria species

CRISPRCasTyper^43^ with default settings was used to analyze CRISPR-Cas loci in all 2712 *Listeria spp*. genomes available from NCBI. The resultant cas_operons.tab output files were parsed to tabulate the number of Type I-B, II-A, II-C, and VI-A loci, and occurrence of associated Cas1 and Cas2 alleles. Type II-A and II-C loci were distinguished by Csn2 family designation.

### Cas9 phylogenetic tree construction

39 diverse type II-A, II-B, and II-C Cas9 protein sequences were aligned with MAFFT using default settings. The resulting alignment was used to generate a statistically-supported phylogenetic tree using the neighbor-joining method in MEGAv11 with default settings and 1000 bootstrap replications.

---

**Figure S1.**
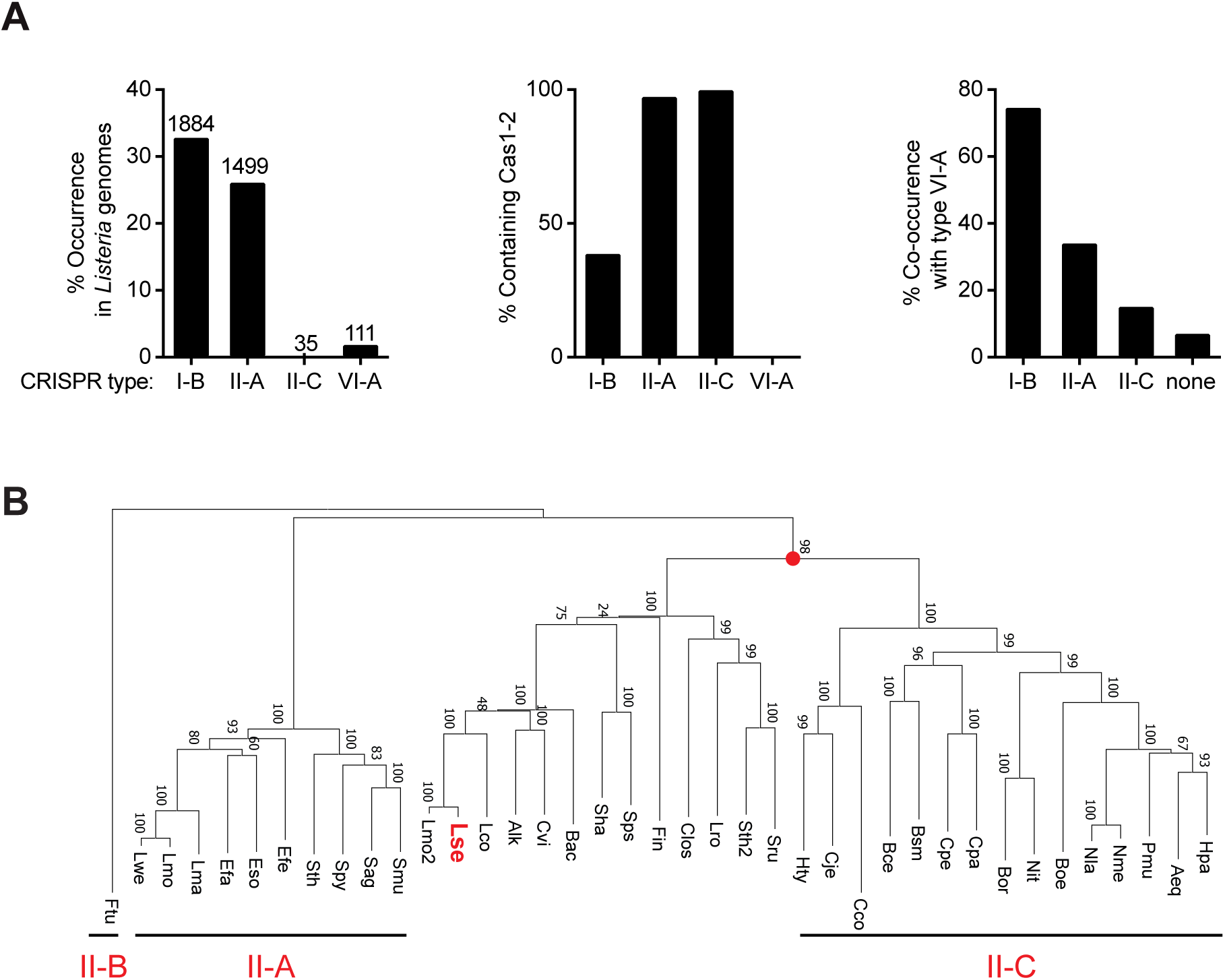
Analysis of CRISPR types and adaptation genes in *Listeria* species. **(A)** 2712 publicly available *Listeria* genomes were analyzed by CRISPRCasTyper to determine the frequency of different CRISPR types (left; number of genomes represented are indicated above the bar), the presence of Cas1 and Cas2 in these CRISPR types (middle), and co-occurrence between type VI-A and other CRISPR types (right). We note that most of the type I-B systems lacking Cas1-2 only lack Cas1, and may have a small (∼240bp) Cas1-like gene that is not detected by CRISPRCasTyper. **(B)** Phylogenetic relationship of 39 diverse Cas9 proteins from type II-A, II-B, and II-C CRISPR-Cas systems. Protein sequences were aligned with MAFFT using default parameters. Phylogenetic tree was constructed using the neighbor-joining method in MEGAv11 with 1000 bootstrap iterations. *L. seeligeri* type II-C Cas9 is indicated in bold red, and clustered with type II-C Cas9 proteins in 98% of iterations.

**Figure S2.**
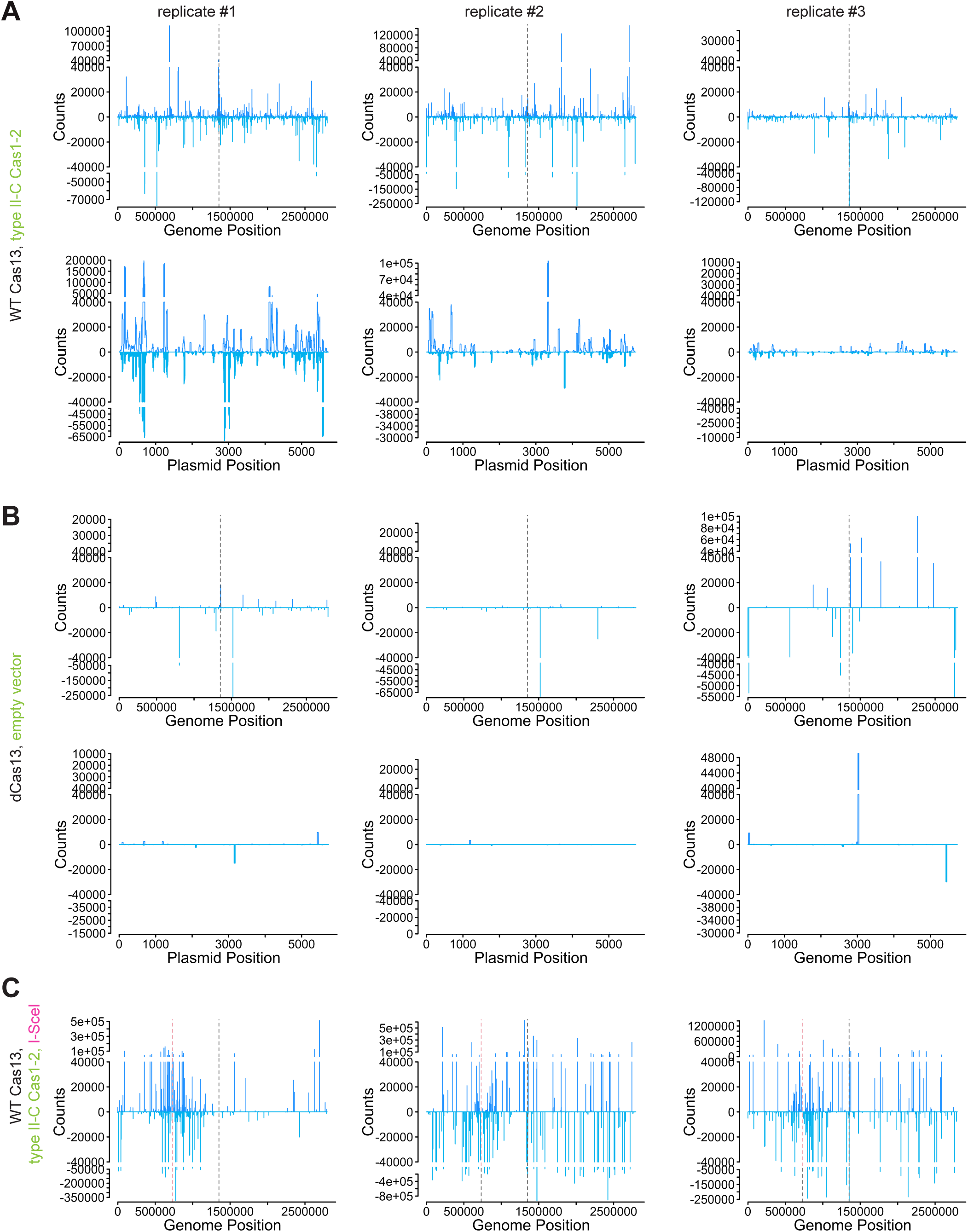
Map of all newly acquired spacers, related to Figure 1. **(A)** Every instance of every spacer mapped to the LS1 genome (top) and plasmid (bottom) for each replicate for the experiment shown in Figure 1E. **(B)** All spacers mapped to the genome or plasmid in the presence of the empty vector control (no Cas1-2). Most sequenced reads are unamplified or PCR artifacts, leading us to conclude that minimal adaptation took place. **(C)** All spacers mapped to the LS1 genome for the experiment shown in Figure 1F.

**Figure S3.**
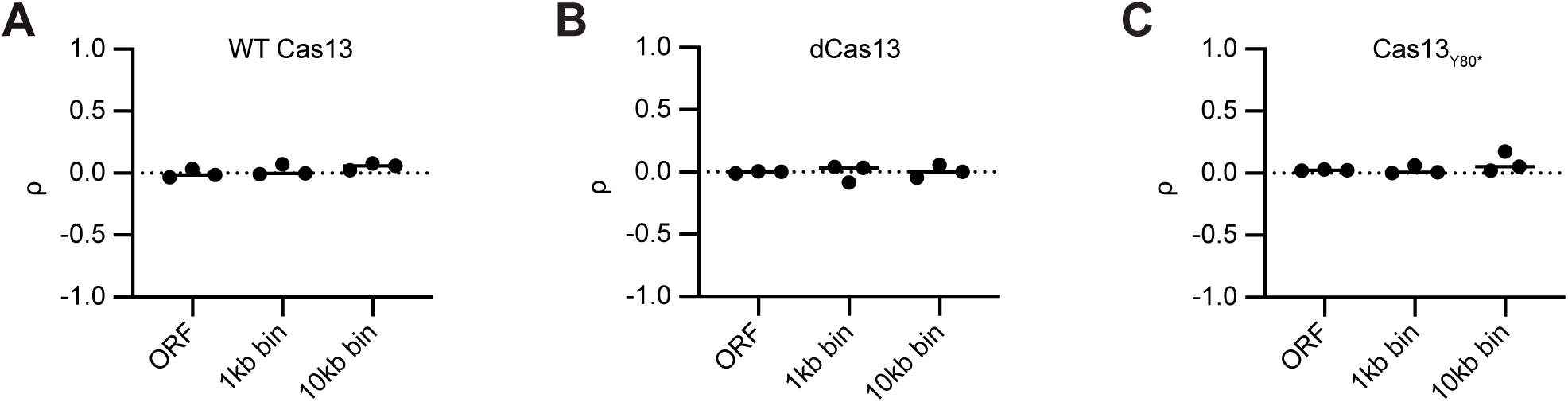
No correlation between spacer acquisition and transcription levels. Frequency of type VI-A spacer acquisition during type II-C induction in an open reading frame (ORF) or bins of specified sizes was compared to transcript levels in that ORF or bin by a Spearman’s rank correlation test. A rho (ρ) value of zero indicated no correlation. **(A)** WT Cas13 samples, related to Figure 1. **(B)** dCas13 samples, related to Figure 2. **(C)** Cas13_Y80*_ samples, related to Figure 2.

**Figure S4.**
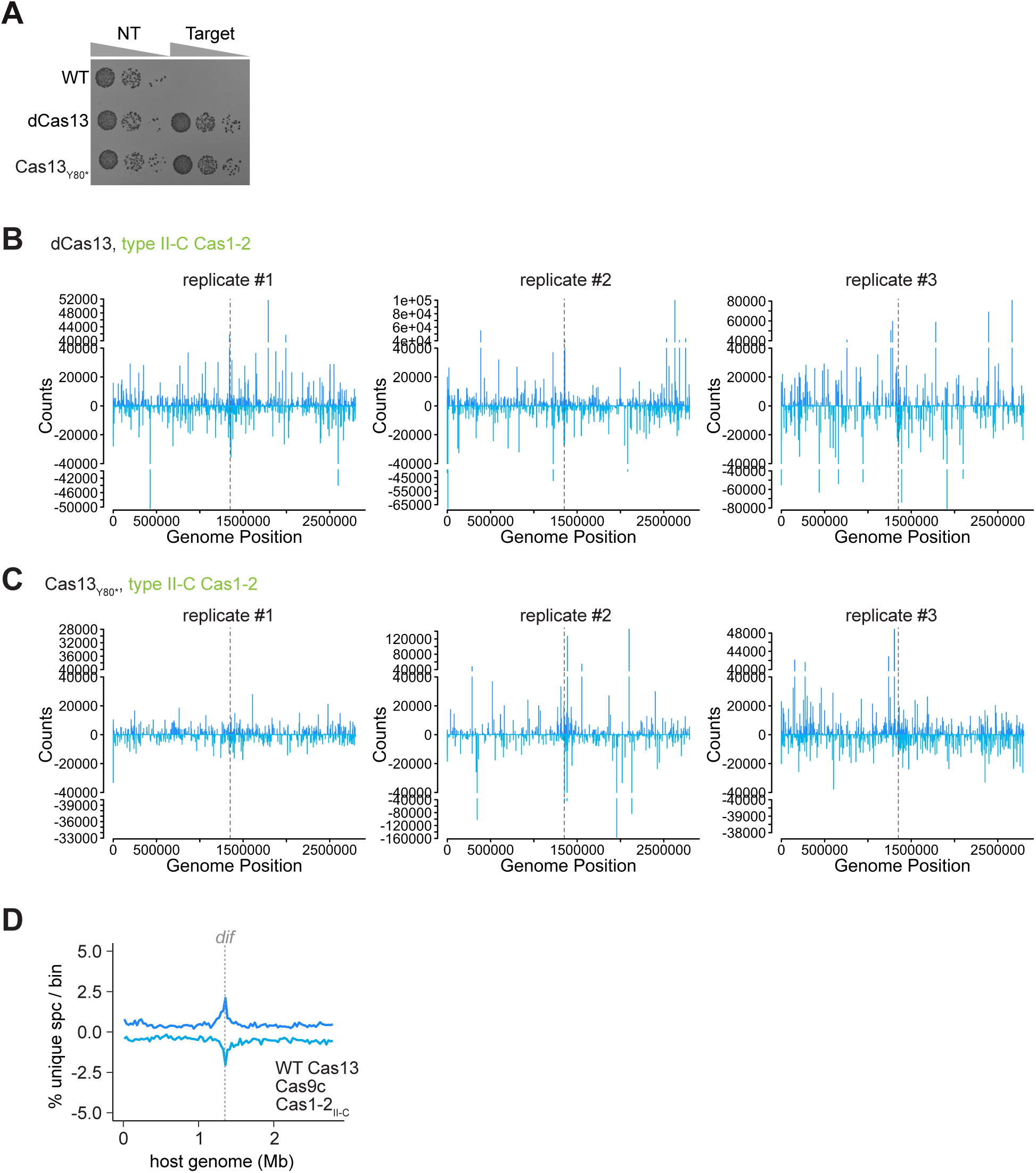
Adaptation is Cas13-independent, related to Figure 2. **(A)** Plasmid challenge assay confirming that dCas13 and Cas13_Y80*_ mutations lead to the loss of immunity. Colony growth indicates tolerance of the Cas13-targeted plasmid. NT= no target control. **(B)** All spacers mapped to the LS1 genome for the experiment shown in Figure 2A with dCas13. **(C)** All spacers mapped to the LS1 genome for the experiment shown in Figure 2B with Cas13_Y80*_. **(D)** Percentage of unique genome-mapped type VI-A spacers acquired per 27976 bp bin during type II-C Cas1-2 induction in the presence of Cas9c. The combination of unique spacers from n=2 replicates is shown. Related to Figure 2E.

**Figure S5.**
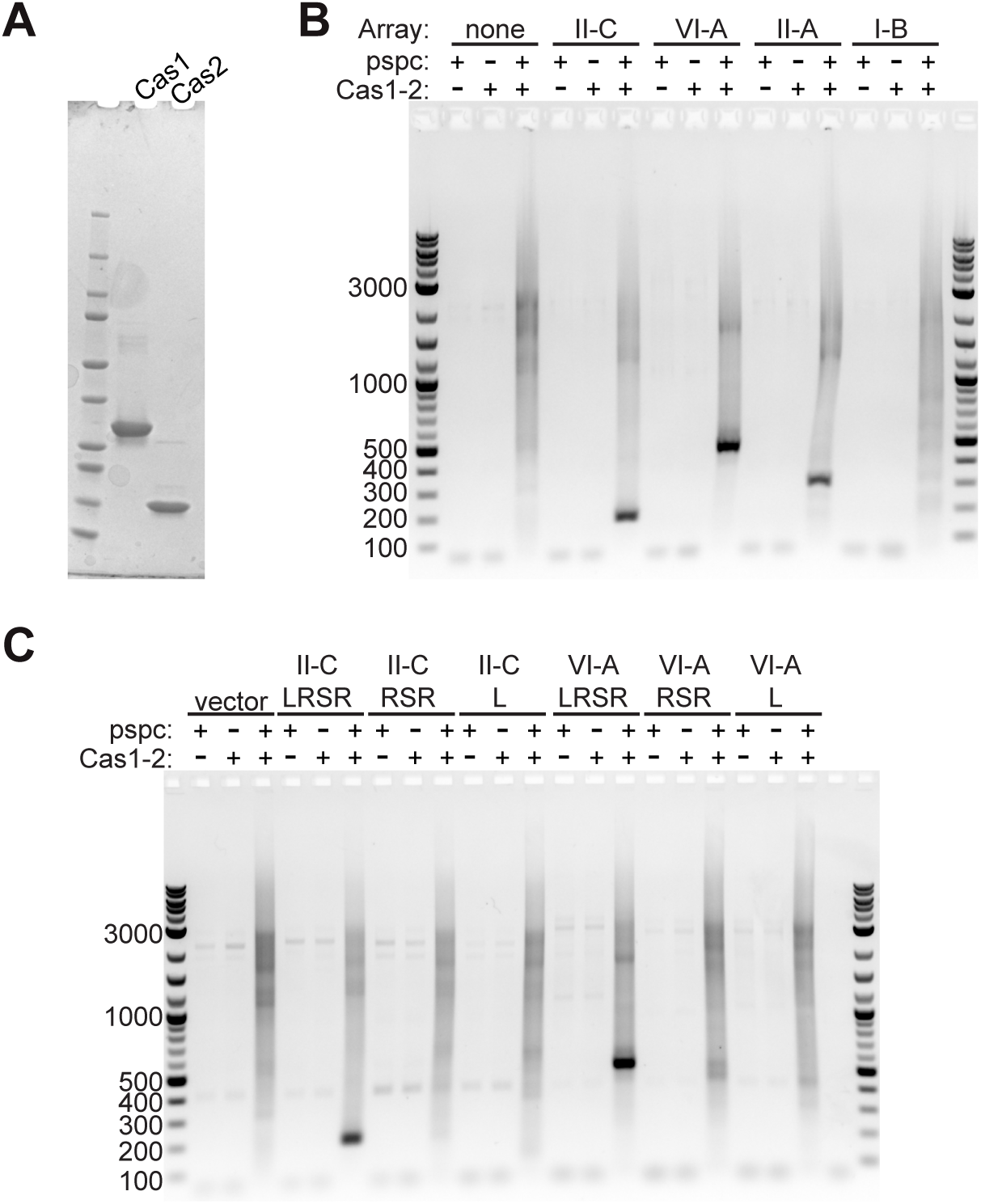
Half-site integration in vitro is dependent on a prespacer and type II-C Cas1-2, related to Figure 3. **(A)** Coomassie stained protein gel of purified Cas1 and Cas2 samples used for in vitro assays. **(B)** Half-site integration PCR experiment shown in Figure 3C with indicated controls. **(C)** Half-site integration PCR experiment shown in Figure 3D with indicated controls. LRSR, Leader-Repeat-Spacer-Repeat. RSR, array only. L, Leader only.

**Figure S6.**
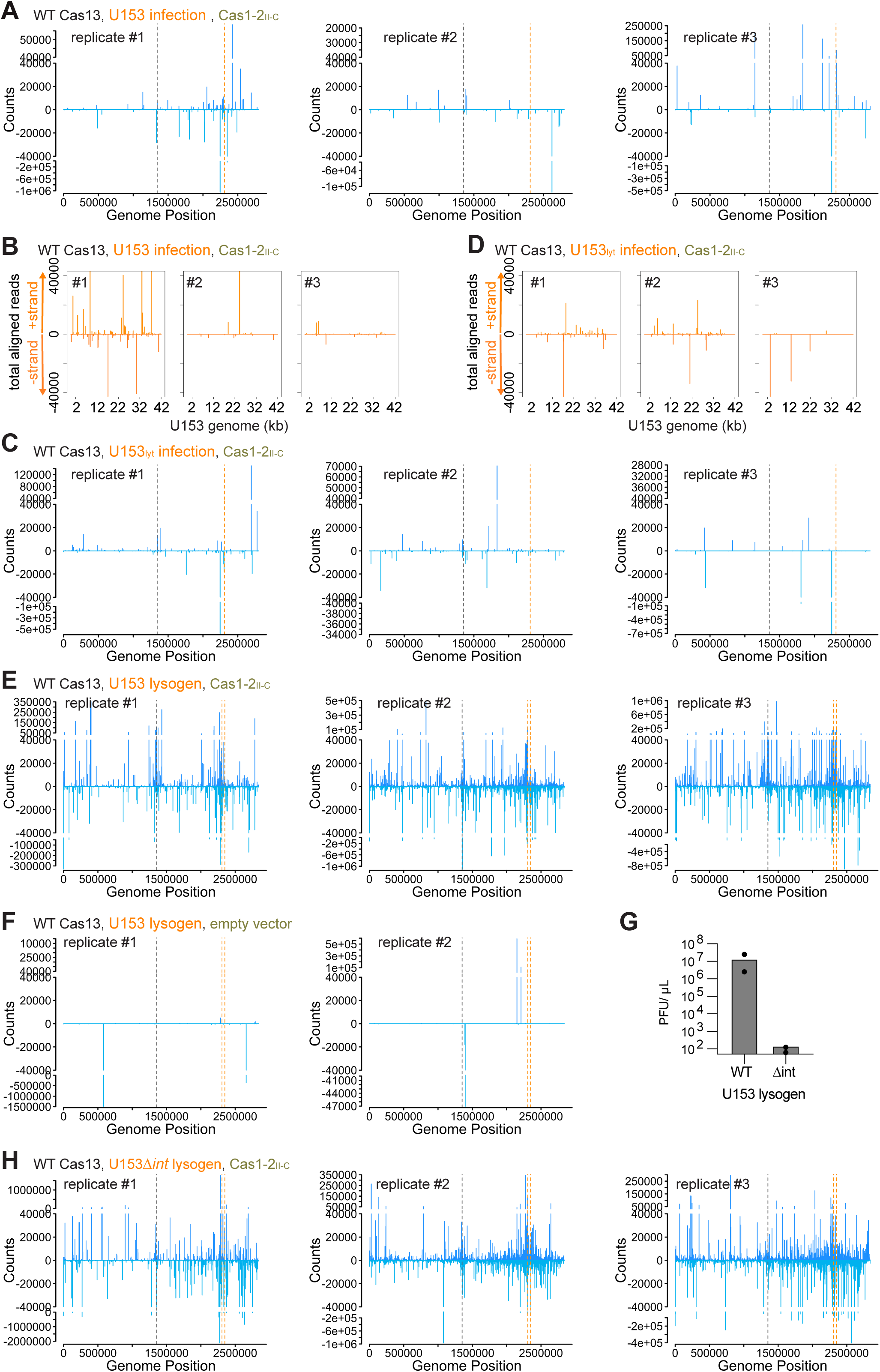
All mapped phage, host, and lysogen targeting spacers; related to Figure 4. **(A)** All spacers mapped to the LS1 genome for the experiment shown in Figure 4C. **(B)** All spacers mapped to the U153 genome for the experiment shown in Figure 4B. **(C)** All spacers mapped to the LS1 genome for the experiment shown in Figure 4E. **(D)** All spacers mapped to the U153 genome for the experiment shown in Figure 4D. **(E)** All spacers mapped to the LS1 U153 lysogen genome for the experiment shown in Figure 4F. **(F)** All spacers mapped to the LS1 U153 lysogen genome in the absence of type II-C Cas1-2 overexpression. **(G)** Plaque forming units produced after overnight induction with mitomycin C from the indicated lysogens. **(H)** All spacers mapped to the LS1 U153 lysogen genome for the experiment shown in Figure 4G.

**Figure S7.**
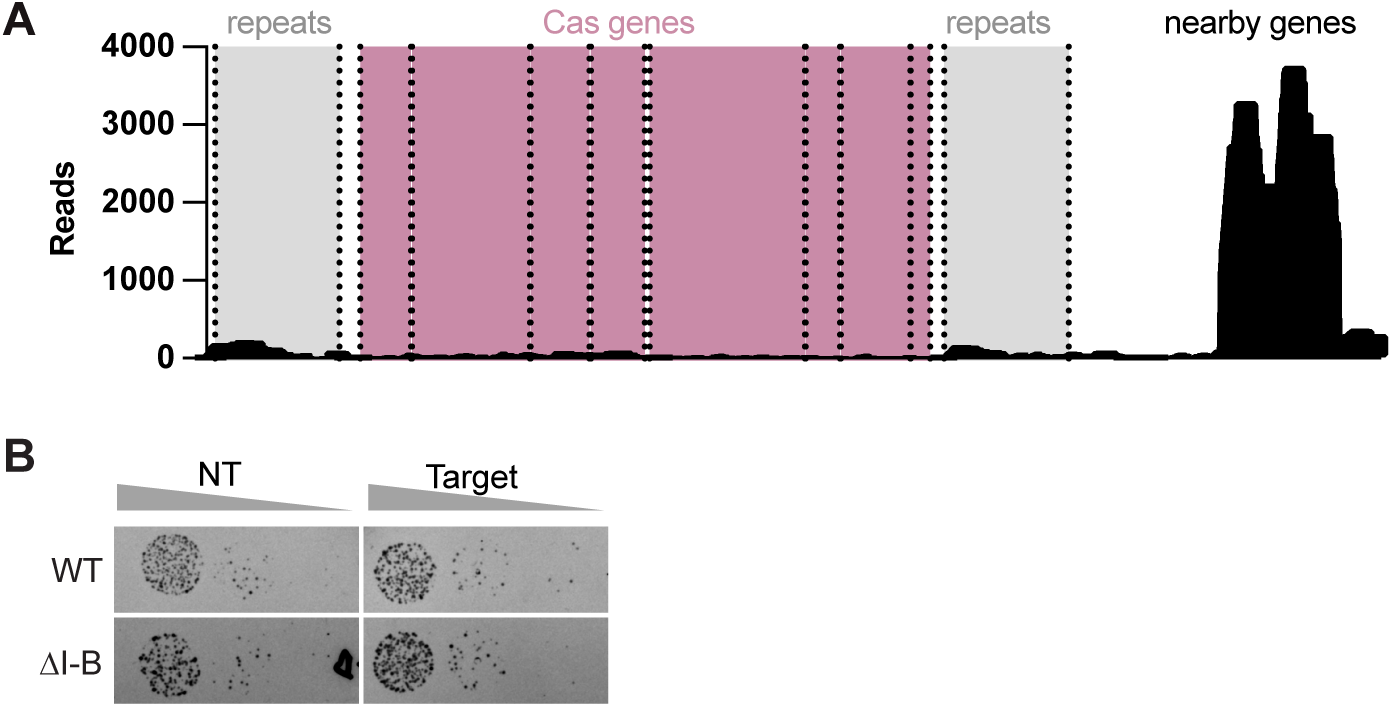
LS1 type I-B CRISPR is not highly expressed or functional in bulk assays. **(A)** Reads from previously published RNA-Seq data showing expression from the LS1 type I-B locus and nearby genes. **(B)** Plasmid challenge assay with a type I-B targeted plasmid or no target vector control (NT) in the presence (WT) or absence (ΔI-B) of the type I-B locus. Despite the presence of the target, there is no restriction of growth. Note that this target is the perfect match of the spacer shown in Figure 5A, which we have previously shown can be functional if the type I-B CRISPR system is expressed (Katz et al. 2024).

**Figure S8.**
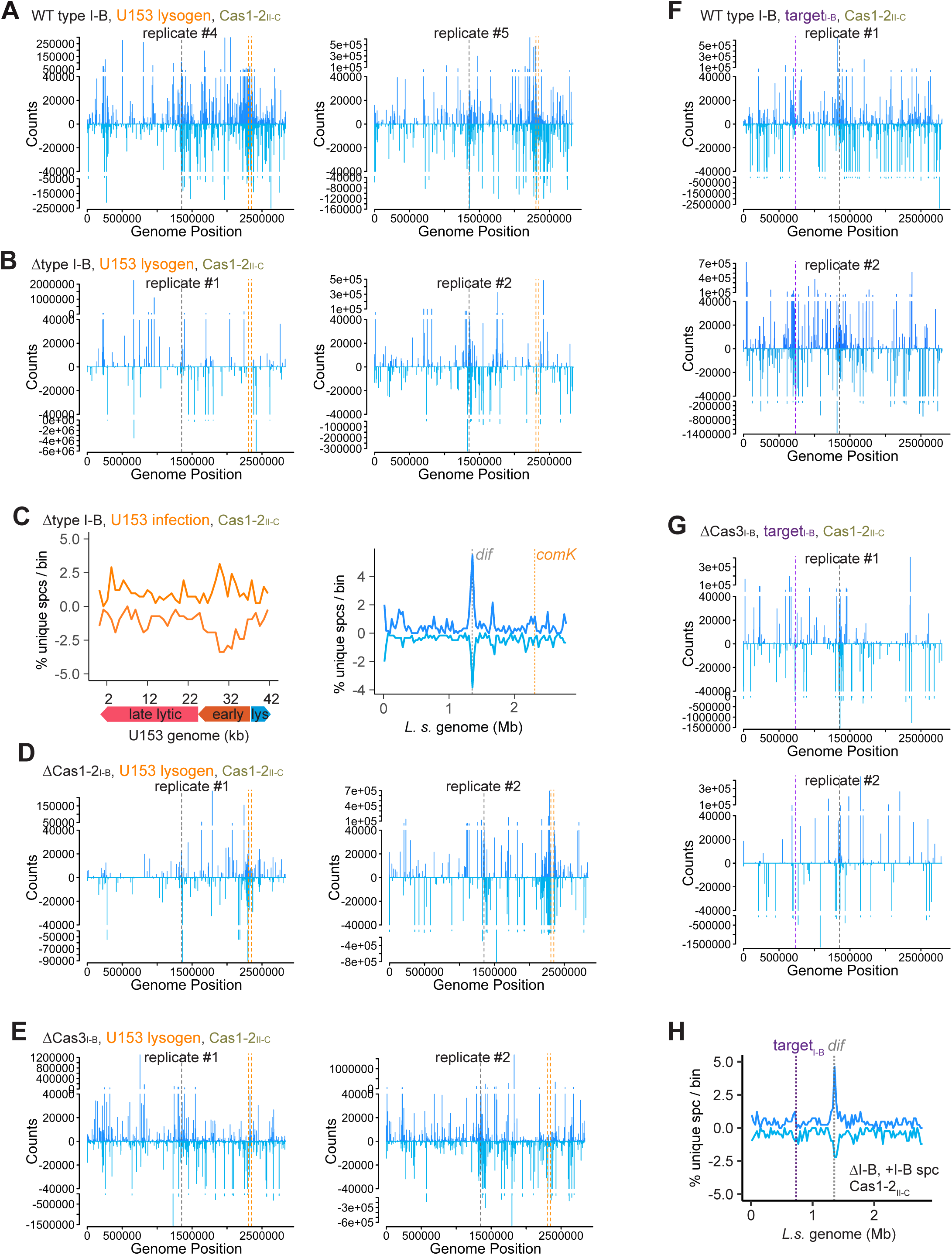
Type I-B CRISPR primes type VI-A spacer acquisition; related to Figure 5. **(A)** All spacers mapped to the LS1 U153 lysogen genome for the experiment shown in Figure 5C. **(B)** All spacers mapped to the LS1 U153 lysogen genome for the experiment shown in Figure 5D. **(C)** Percentage of unique spacers per bin (using the same bin sizes as Figure 4) acquired during concomitant type II-C Cas1-2 induction and U153 infection of ΔI-B, mapped to the phage or host genome. The combination of unique spacers from n=2 replicates is shown. **(D)** All spacers mapped to the LS1 U153 lysogen genome for the experiment shown in Figure 5E. **(E)** All spacers mapped to the LS1 U153 lysogen genome for the experiment shown in Figure 5F. **(F)** All spacers mapped to the LS1 genome for the experiment shown in Figure 5G. **(G)** All spacers mapped to the LS1 genome for the experiment shown in Figure 5H. **(H)** Percentage of unique genome-mapped spacers acquired per 1% bin during type II-C Cas1-2 induction in ΔI-B in the presence of a type I-B spacer targeting the genomic location noted by the gray line. The type I-B spacer is under control of the native promoter. Results are from a single replicate.

